# Pangenome and comparative genomics analysis reveal patterns of quorum sensing and antimicrobial resistance in *Salmonella* Lineages from African Food Chains

**DOI:** 10.1101/2025.07.28.667100

**Authors:** M. N. Muigano, F. M. Musila, Z. Li

## Abstract

*Salmonella enterica* is a major public health concern in Africa because of its increasing antimicrobial resistance (AMR) to multiple antibiotics that threatens effective treatment and control. In this study, we analyzed 135 publicly available antimicrobial resistant Salmonella genomes from African food sources. Using a pan genomic analysis approach, we identified a pangenome of 15,269 genes, including a core genome of ∼3.5 Mbp with 3,350 core genes (99-100% prevalence) and a substantial accessory genome comprising 183 soft-core, 1,441 shell, and 10,295 cloud genes. We found substantial genomic diversity among the AMR-positive *Salmonella enterica* isolates. AMR profiling revealed widespread distribution of resistance genes, with *aac(6’)-laa_1* present in nearly all genomes, and *tet(A)*, *sul2*, and *fosA71* frequently co-occurring. Core virulence genes such as *invA*, *ssaL*, and *steA* were nearly universally conserved, suggesting essential roles in pathogenesis. Sporadic rare virulence genes like *papE*, *tcpC*, and *ybtP* were present in the genome pointing to lineage-specific patterns. The presence of macrolide inactivation genes (*ereA*) points to a huge risk of resistance to the globally important antibiotic Azithromycin. Quorum-sensing genes, including the AI-2 system (*luxS*, *lsrA-D*, *lsrK*) and biofilm formation genes were widely conserved across the isolates, highlighting their potential role in adaptation and the spread of antimicrobial resistance in foodborne *Salmonella* in Africa. Hierarchical clustering based on core genome multilocus sequence typing (cgMLST) revealed monophyletic clades for major serovars like *Enteritidis* and *Infantis*. Shannon diversity indices revealed geographical variation in AMR gene richness, with Burkina Faso, South Africa and Tunisia showing high intra-country diversity. Our analysis identified lineage-associated accessory genes, hence providing potential markers for surveillance. This study offers valuable insights into the genomic architecture and AMR landscape of *Salmonella* in Africa and underscores the need for expanded genomic surveillance across diverse geographic and food production contexts.

## 1. Introduction

*Salmonella* infections are a major public health concern in the world and is implicated in a range of infections affecting both humans and animals. Every year, between 200 million and 1 billion *Salmonella* infections are reported around the world, with majority of them being foodborne-related (Lamichhane et al., 2024). Additionally, some estimates put cases of *Salmonella*-related deaths at over 150,000 (Lamichhane et al., 2024; Nazir et al., 2025). The economic burden is equally huge, with conservative estimates from 2012 putting the figure at $3.3 billion (Hoffmann et al., 2012). Several of its serovars are known for their zoonotic potential, with foodborne transmission representing a major route of infection.

The risk of *Salmonella* infection is higher in low-income countries such as those in Africa where hygiene and sanitation conditions may be inadequate (Ngogo et al., 2020). In many parts of the world including Africa, foodborne salmonellosis is made worse by the rising prevalence of antimicrobial-resistant (AMR) strains circulating in the food chain (Tack et al., 2020). The frequent recovery of AMR *Salmonella* from animal-derived food products has complicated treatment and control strategies (Salam et al., 2023). The problem is likely worse in resource-limited settings like those in many African countries.

AMR in *Salmonella* is often driven by a combination of mobile genetic elements such as plasmids, transposons, integrons, and resistance islands, alongside chromosomal mutations (Algarni et al., 2022; Partridge et al., 2018). These elements not only enable survival in antibiotic-exposed environments but also support the horizontal and vertical transfer of resistance genes across bacterial species (Partridge et al., 2018). In addition, *Salmonella* species have intrinsic mechanisms for resisting antimicrobial therapy (Chaudhari et al., 2023). The presence of AMR genes in foodborne isolates suggests that these strains may act as important reservoirs of resistance, with implications for both animal and human health.

An analysis of large datasets of *Salmonella* genomes may offer useful insights into the AMR landscape. Advances in comparative genomics have made it possible to explore genetic diversity across bacterial populations in greater detail. One such approach is pangenome analysis, which allows researchers to examine both the conserved (core) and variable (accessory) gene content across multiple strains (Khan et al., 2023). This can offer insights into traits related to host adaptation, virulence, and antimicrobial resistance. Accessory genes, in particular, may include lineage-specific elements or mobile genetic factors that go undetected in single-genome analyses (Costa et al., 2020). Thus, in this study, we performed a pangenome analysis of AMR-positive *Salmonella enterica* strains isolated from food sources across different African regions. Publicly available genome assemblies were used to identify patterns in core and accessory gene content and to explore possible associations with antimicrobial resistance. By comparing strains from varied geographic locations, we aimed to uncover genetic markers or signatures that may be linked to resistance phenotypes. Our findings contribute to the growing body of genomic data on African *Salmonella* isolates and may inform future efforts in surveillance, diagnostics, and the management of foodborne infections.

## 2. Materials and Methods

### 2.1 Data collection and genome retrieval

In this study we retrieved a total of 526 *Salmonella enterica* genome assemblies from EnteroBase, a publicly accessible bacterial genome database and analysis platform (https://enterobase.warwick.ac.uk/). The dataset was filtered to retain isolates of African origin, recovered exclusively from food-associated sources, including meat, milk, eggs, vegetables, and related processing environments (https://enterobase.warwick.ac.uk/species/senterica/search_strains?query=workspace:139979). To minimize the presence of temporal outliers, we restricted our search to assemblies collected after 2000. The search was conducted on 05 July 2025. To further refine the dataset, genome metadata were examined for antimicrobial resistance (AMR) profiles, resulting in the selection of 135 AMR-positive *Salmonella* genomes for downstream comparative analysis. Information regarding the isolates, including geographic distribution, food source, sequence type (ST), and inferred serovar, was extracted from EnteroBase records and are available in Supplementary Table 1.

### 2.2. Core Genome Multilocus Sequence Analysis

We used EnteroBase’s in-built core genome multilocus sequence typing (cgMLST) scheme for initial high-resolution genotyping of the isolates (Accessed on July 07, 2025). We assigned clonal groups to the isolates using the Hierarchical Clustering of Core Genome Multi-Locus Sequence Typing (HierCC-cgMLST). We constructed a GrapeTree using the NINJA algorithm to visualize the phylogenetic structure and clustering of the strains based on allelic differences across core loci for all the 526 assemblies. This allowed for the preliminary identification of lineage patterns and inter-strain relatedness prior to whole-genome analysis.

### 2.3 Genome annotation and quality control

We downloaded the genome assemblies corresponding to the selected 135 AMR-positive isolates in FASTA format for further processing and annotation. The downloaded genome assemblies were subjected to structural and functional annotation using the Prokka v1.14.6 pipeline (Seemann, 2014), within a Linux-based Ubuntu environment. Each genome was processed individually with Prokka’s default bacterial gene prediction settings, and the output comprised GenBank (GBK), GFF3, and protein FASTA (FAA) files for each strain. The annotation workflow was implemented using a custom Bash script within a dedicated Conda environment. To ensure annotation consistency and compatibility with downstream pangenome analysis, all FASTA headers were pre-processed to remove complex characters and standardize sequence identifiers. Quality control of annotation outputs was confirmed by examining the number of annotated coding sequences (CDS), rRNA and tRNA genes, and checking for the presence of complete GFF files. A total of 135 GFF files were successfully generated and utilized for comparative genomic analysis.

### 2.4 Pangenome construction and core genome alignment

Comparative pangenome analysis was performed using the Roary pipeline (v3.13.0), a rapid large-scale tool for prokaryotic pangenome construction (Page et al., 2015). All 135 annotated .gff files were provided as input to Roary. The analysis was executed with the -e and --mafft options enabled to ensure accurate core gene alignment using MAFFT. The clustering identity threshold was set to 95%, and genes present in at least 99% of genomes were defined as part of the soft-core genome. Output files included the core gene alignment file (core_gene_alignment.aln), gene presence/absence matrix (gene_presence_absence.csv), and a variety of cluster statistics and supporting data.

### 2.5 Serovar Prediction and Phylogenetic reconstruction

Assemblies with unknown serovar composition within the AMR-positive *Salmonella* genomes were assigned using the *Salmonella* In Silico Typing Resource (SISTR) (Yoshida et al., 2016). SISTR integrates antigenic typing and core genome MLST (cgMLST) to assign serovar predictions. In our study, we analyzed the cleaned genome assemblies using the command-line version of SISTR. The output included key metadata such as antigenic formula, serovar designation, and cgMLST information. Predicted serovars were then integrated with the phylogenetic tree and visualized in iTOL using color strip annotations, with rare serovars grouped under the category “Other” to improve interpretability.

Phylogenetic inference of the 135 *Salmonella* genomes was performed using the aligned core genome using FastTree (v2.1.11) (Price et al., 2010). The resulting phylogenetic tree was generated in Newick format and visualized using Interactive Tree of Life (iTOL v5) for downstream interpretation (Letunic and Bork, 2021). The tree topology was further integrated with metadata, including country of origin, ST, and AMR class, to explore potential associations between phylogeny and phenotypic traits.

### 2.6 Evaluating Genomic Plasticity with Heap’s Law Fitting

To assess the openness of the pangenome, we applied Heap’s Law, a power-law model originally adapted to microbial genomics by Tettelin and co-authors (Tettelin et al., 2008). This approach models the cumulative number of unique genes (P) discovered as more genomes (N) are added, using the equation

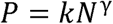

where k and γ are empirical constants derived through nonlinear curve fitting. The exponent γ serves as an indicator of pangenome dynamics with values closer to 1 implying high genomic fluidity and openness, while values near 0 suggest saturation and a closed gene repertoire. We used the criteria proposed by Hyun et al. (2022) to classify pangenomes as follows: γ < 0.3 (closed) and γ > 0.3 (intermediate to open).

### 2.7 Detection and analysis of antimicrobial resistance, plasmid, and virulence genes

To investigate the genetic basis of antimicrobial resistance, we screened all genome assemblies for acquired AMR genes using the ABRicate tool (v1.0.1) (https://github.com/tseemann/abricate) with the ResFinder database (Zankari et al., 2012). The search parameters included a minimum identity of 90% and minimum coverage of 80%. Detected genes were grouped according to resistance class (e.g., beta-lactams, aminoglycosides, fluoroquinolones, sulfonamides), and their distribution was cross-referenced with the accessory genome clusters obtained from Roary to identify potential resistance-associated genes and lineage-specific markers. The presence of resistance determinants was also evaluated in relation to phylogenetic clades derived from core genome alignment. Virulence-associated genes were identified using the VFDB database in ABRicate (Chen et al., 2016), while plasmid replicons were detected using the PlasmidFinder database (Carattoli et al., 2014).

### 2.8 Data analysis and visualization

Post-processing and downstream analyses were conducted in Python 3.10 using packages like pandas, seaborn, matplotlib, and scikit-learn. The gene presence/absence matrix generated by Roary (gene_presence_absence.csv) was used as input for exploratory analysis of genome-wide gene content variation (See Supplementary File 2). Heatmaps and binary matrices were constructed to visualize the distribution of accessory genes across isolates. Core genome phylogeny generated by FastTree was visualized using iTOL (Interactive Tree of Life) with metadata layers including country of origin, ST, and AMR class. The alignment file (core_gene_alignment.aln) was used as input to infer relationships among the strains and assess potential clustering by AMR gene profiles or food source. Additionally, presence/absence patterns of acquired AMR genes (as identified by ABRicate + ResFinder) were mapped to Roary accessory gene clusters to identify candidate lineage-specific or resistance-associated markers. Genes unique to AMR-positive isolates or clades were shortlisted for further investigation. Data curation, figure generation, and final visualization were performed in Jupyter Notebook.

## 3. Results

### 3.1 Hierarchical Core Genome Phylogeny and Serovar-Level Clustering of *Salmonella* Isolates

To investigate the population structure of *Salmonella enterica* isolates from Africa, we analyzed phylogenetic relationships using cgMLST-based hierarchical clustering (HierCC) at two levels of resolution, along with *in silico* serotyping from SISTR1 (Figure 1). At the fine-scale HC100 level, closely related isolates clustered tightly, indicating close genetic similarity. Broader clustering at the HC2000 level was evident with distinct super-lineages encompassing multiple serovars. The SISTR1-based tree identified the most prevalent serovars as *Typhimurium* (n=38), *Enteritidis* (n=34), Kentucky (n=21), and Newport (n=30). Meanwhile, HC2000 clustering revealed dominance of super-lineages 2 (n=60), 12 (n=39), and 724 (n=27), whereas HC100 distinguished finer clades such as 724 (n=26), 924 (n=25), and 48 (n=22). Serovars like *Enteritidis* and *Infantis* formed tight monophyletic clusters across all methods, supporting clonal expansion. In contrast, serovars with high counts but spread across multiple HC100 clusters, such as *Typhimurium* and Newport, showed greater intra-serovar diversity. Furthermore, the grouping of diverse serovars within single HC2000 lineages highlights the presence of broader ancestral lineages circulating across the continent.

**Figure 1:**
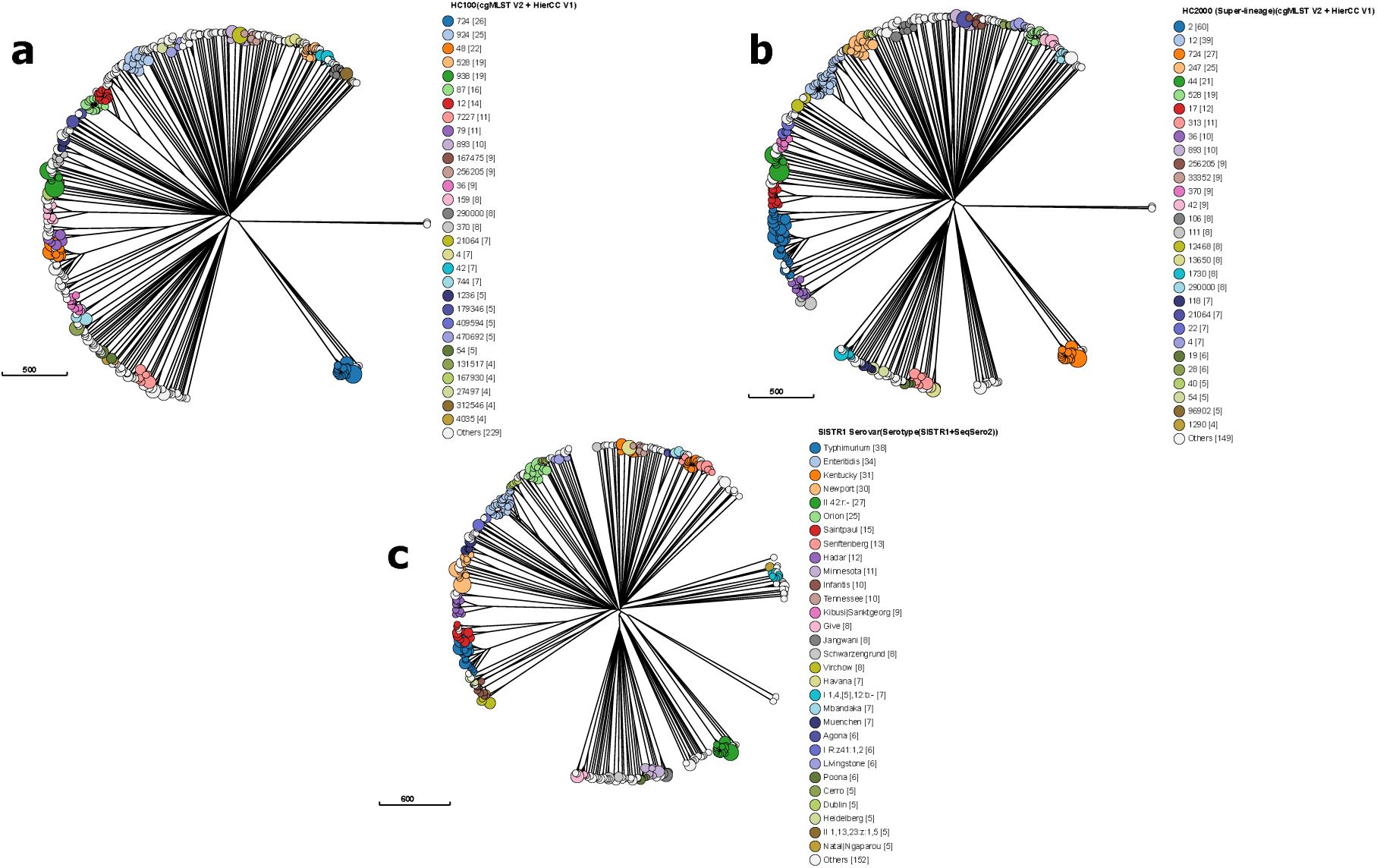
MLST structure of Salmonella in African food chains in Enterobase showing HC100 (cgMLST + HierCC) clustering (A), super-lineage hierarchical clustering (B), and SISTR serovar typing (C). Each node represents a single ST and those with the same color belong to the same serovar. GrapTree clustering was done with the NINJA NJ algorithm.

A total of 32 distinct *Salmonella* serovars were identified among the 135 AMR-positive genomes using the SISTR typing platform (Figure 2A). These serovars represented a wide range of antigenic profiles and genetic diversity within the dataset. Key serovars included *Kentucky*, *Enteritidis*, *Minnesota*, *Muenster*, *Schwarzengrund*, and *Newport*, with *Kentucky* and its predicted variants constituting a majority of the isolates. Several rare or infrequently observed serovars were also detected but were grouped under the category “Other” for visualization clarity in the phylogenetic tree. This diversity suggests the presence of multiple evolutionary lineages and may indicate differing ecological niches or sources of infection. Figure 2A below shows an overlay of these serovar classifications onto the country and source type metadata. To complement the phylogenetic analysis and further explore genomic diversity, a Phandango plot was generated to visualize the distribution of core and accessory genes across the Salmonella genomes (Figure 2B). The Phandango plot shows the patterns of conserved and variable genes within and across serovars. The heatmap shows strong gene presence from 0–1.3 kb, which corresponds to the core genome. These are the genes conserved across nearly all strains. Beyond 1.3 kb, the intensity of gene presence decreases toward 6.4 kb, indicating accessory genes that are present in only a subset of strains. This trend is reflected in the line graph beneath the heatmap, which depicts the proportion of strains harboring each gene. The line remains near 100% within the core genome region, then progressively declines across the accessory genome, ultimately approaching zero. Overall, this plot highlights the distinction between conserved housekeeping genes and variable, strain-specific genes, which may be associated with adaptation, virulence, or environmental specificity.

**Figure 2:**
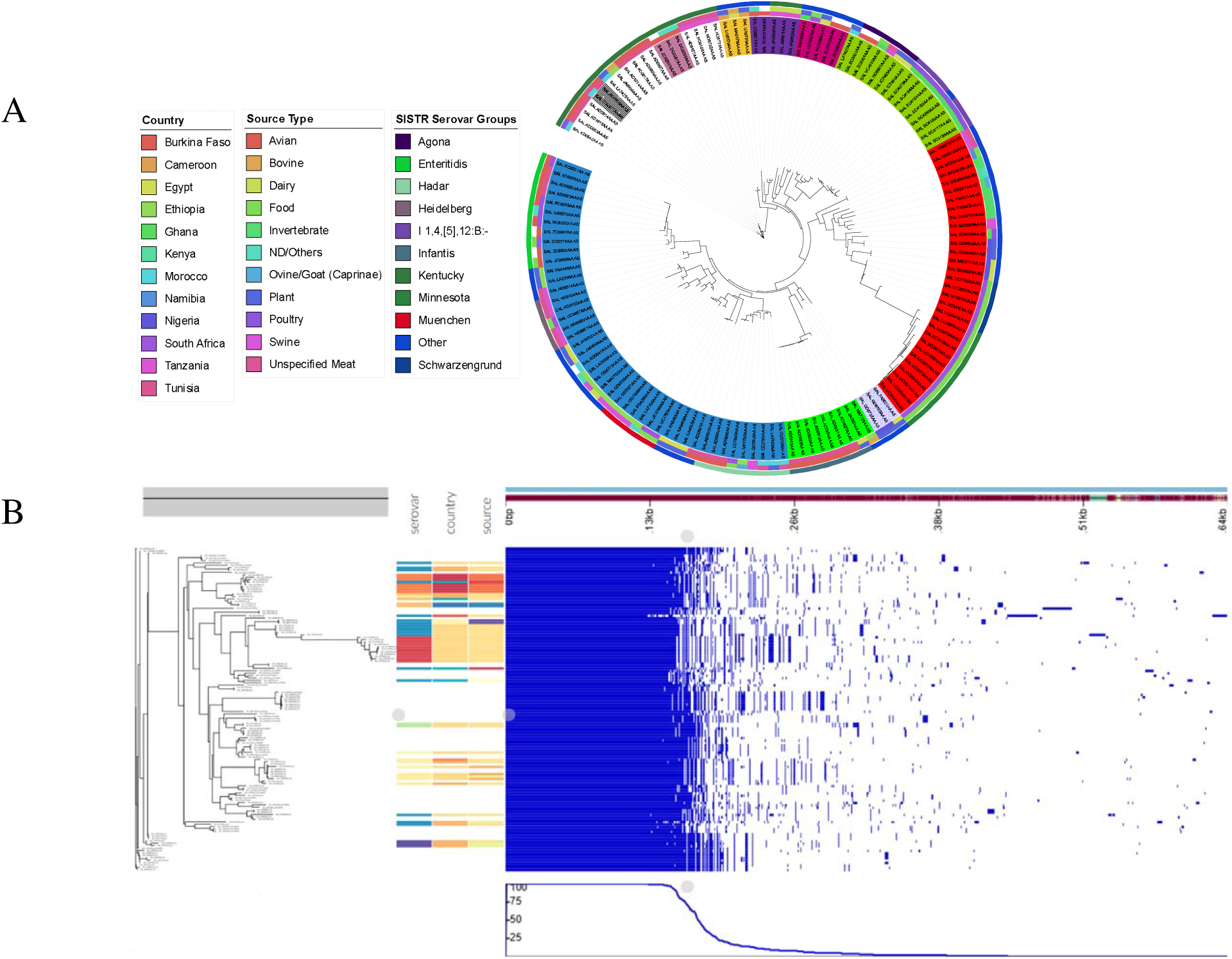
Core genome phylogeny of 135 AMR-positive *Salmonella enterica* isolates from African food chains (A) and a corresponding Phandango plot (B). The phylogenetic tree was constructed on iTOL using cgMLST-based allele profiles. Metadata rings represent serovar (inner ring), source type (middle ring), and country of isolation (outer ring). The Phandango plot shows the presence (blue) and absence (white) of genes in the core genome of 135 AMR positive *Salmonella* isolates.

### 3.2 Pangenome analysis

Pan-genome analysis of 135 *Salmonella enterica* isolates from African food chains revealed a total of 15,269 unique gene clusters with an average gene length of 1,054.62 bp. Of these, 3,350 genes (21.9%), equivalent to ∼3.53 Mbp, were classified as core genes as they were present in 99–100% of all genomes. This suggests a conserved set of functions essential across the *Salmonella* lineages in the African samples. An additional 183 genes (1.2%), which were found in 95–<99% of the isolates were identified as soft-core genes (Figure 3). The accessory genome comprised the majority of the gene clusters. It included 1,441 genes (9.4%) that formed the shell genome and which was shared by 15–<95% of isolates. The accessory genome also included a large proportion (10,295 genes; 67.4%) that fell within the cloud genome, and which were present in fewer than 15% of the genomes. This extensive cloud genome highlights substantial genetic diversity and plasticity across the sampled isolates. This could be due to lineage-specific adaptations, horizontal gene transfer events, and differential environmental pressures across the continent.

**Figure 3:**
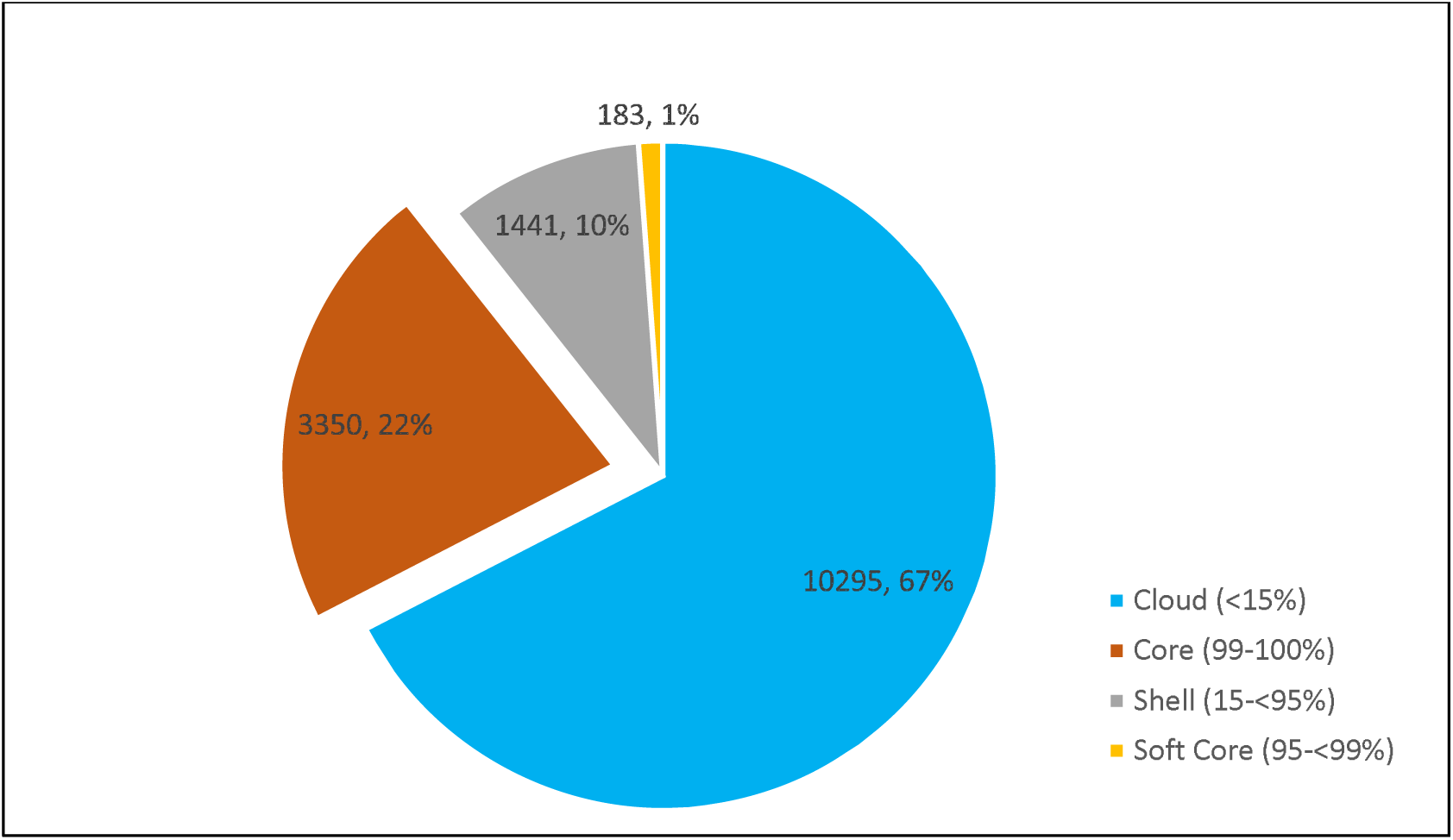
The distribution of genes found in the genomes of AMR *Salmonella* strains from food chains in Africa

The pan-genome curve shown in Figure 4 exhibited a steady increase in the cumulative number of genes as new *Salmonella* genomes were added, indicative of continuous gene acquisition across isolates. The growth curve showed a characteristic upward trajectory that began to gradually plateau as more genomes were included, suggesting that the majority of accessory genes had been captured within the dataset. Although the general trend was smooth, occasional minor sharp rises were observed, likely representing rare strain-specific or niche-associated genes introduced by a few individual genomes. The overall pattern of the curve aligns with an intermediate to closed pan-genome, as further supported by Heap’s Law fitting (Tettelin et al., 2008). The fitted Heap’s Law model yielded an exponent (γ) of 0.2844, indicating a moderately closed pan-genome. This suggests that the *Salmonella* genome from African food chains has little the capacity to acquire novel genes as additional genomes are sequenced. While new genes continue to appear with additional genomes, the rate of gene discovery diminishes over time, which may be indicative of a reflection of a semi-diverse but not unlimited gene pool. Such a trend is characteristic of bacterial populations that engage in limited horizontal gene transfer and occupy ecologically overlapping environments, such as food chains across the African continent. The core-genome curve displayed a progressive decline in the number of conserved genes shared across all genomes as the number of *Salmonella* assemblies increased. This trend reflects the expected reduction in universally shared genes as genomic diversity expands. The curve gradually flattened after approximately 65 genomes and this could be an indication of a stabilization of the core genome size. A single minor sharp drop was observed around that point, suggesting the inclusion of a genomically divergent strain that lacked a few otherwise conserved genes. This pattern is typical of bacterial populations with a relatively stable and conserved core genome, where most essential housekeeping genes are retained across strains. The eventual plateau indicates that adding more genomes is unlikely to significantly reduce the core gene set, an observation that is consistent with a closed or nearly closed core genome architecture. So, functional constraints on essential genes across food-associated *Salmonella* strains may exist and implies limited variability in the core functions necessary for survival and pathogenicity.

**Figure 4:**
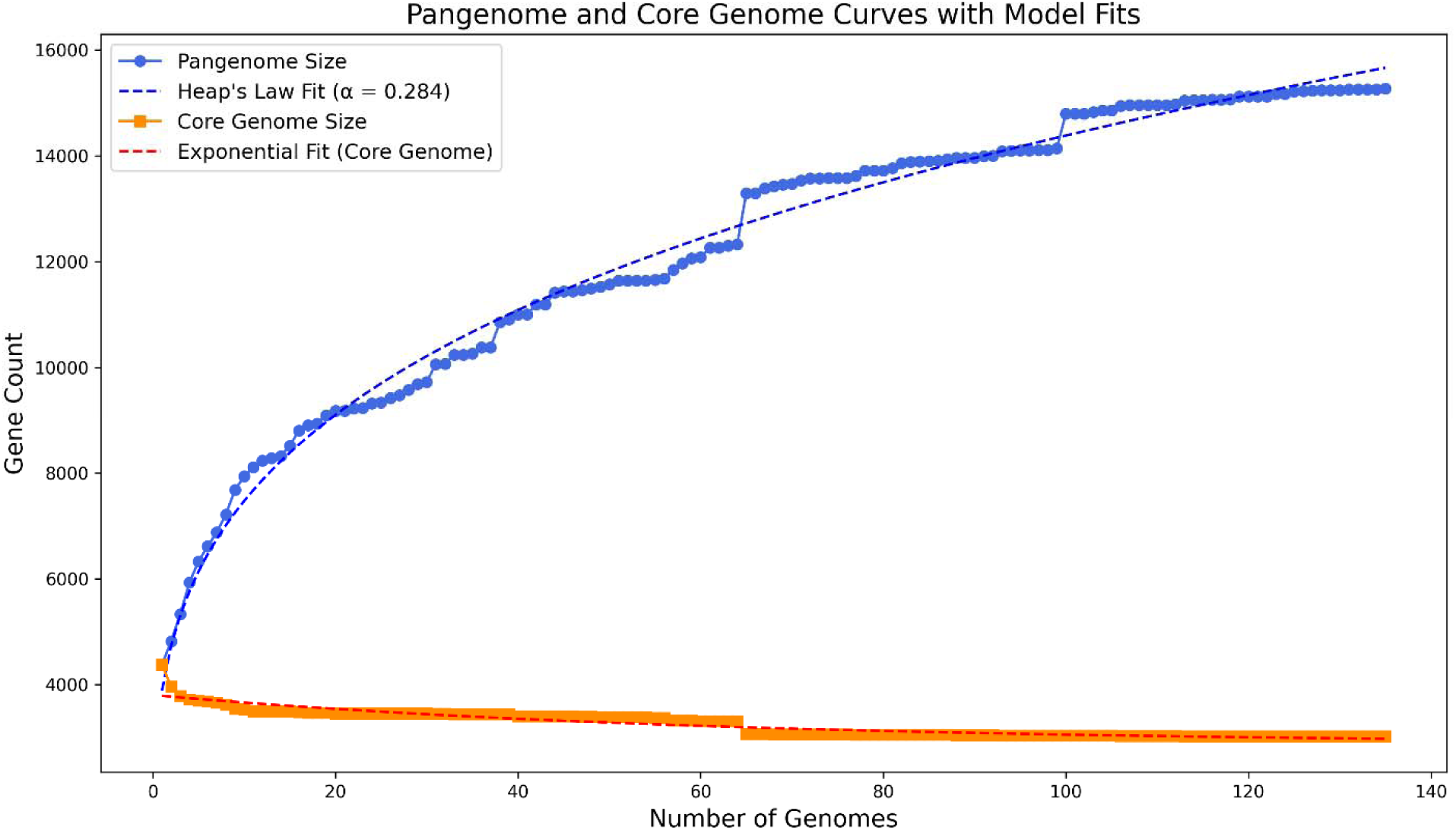
Pan-genome expansion dynamics of 135 AMR-positive *Salmonella enterica* isolates. The pan-genome accumulation curve (royal blue) shows the cumulative number of genes as genomes are sequentially added. Heap’s Law model fit to the pan-genome (dotted blue line) illustrates an open pan-genome architecture (γ < 1), suggestive of ongoing horizontal gene acquisition and high genomic plasticity within the population. Core genome contraction curve (dark orange) of AMR-positive *Salmonella enterica* isolates from African food chains shows the decreasing trend in the number of shared (core) genes as more genomes are added. This indicates an increase in genetic diversity among the isolates. The plateauing pattern suggests a conserved core set of genes essential for the survival and maintenance of fundamental biological functions.

Figure 5 shows the gene frequency distribution of the 135 AMR-positive *Salmonella* genomes with a characteristic U-shaped profile, consistent with a dynamic bacterial pangenome. A prominent peak was observed at the far left of the histogram, indicating a high number of unique genes (n = 4,649) found in only one genome. These strain-specific genes may represent mobile genetic elements, recent horizontal acquisitions, or adaptations to niche-specific environments. As the number of genomes sharing a gene increased, the frequency of such genes decreased steadily, flattening off around the 40-genome mark, reflecting a large set of accessory genes (n = 7,604) that are variably distributed across the population. A second peak emerged at the far right of the histogram, representing core genes (n = 3,016) that are conserved across all 135 genomes. This bimodal distribution emphasizes the presence of both a stable, conserved genomic backbone and a flexible gene pool. This could point to the semi-closed nature of the *Salmonella* pangenome in AMR-associated isolates from African food chains.

**Figure 5:**
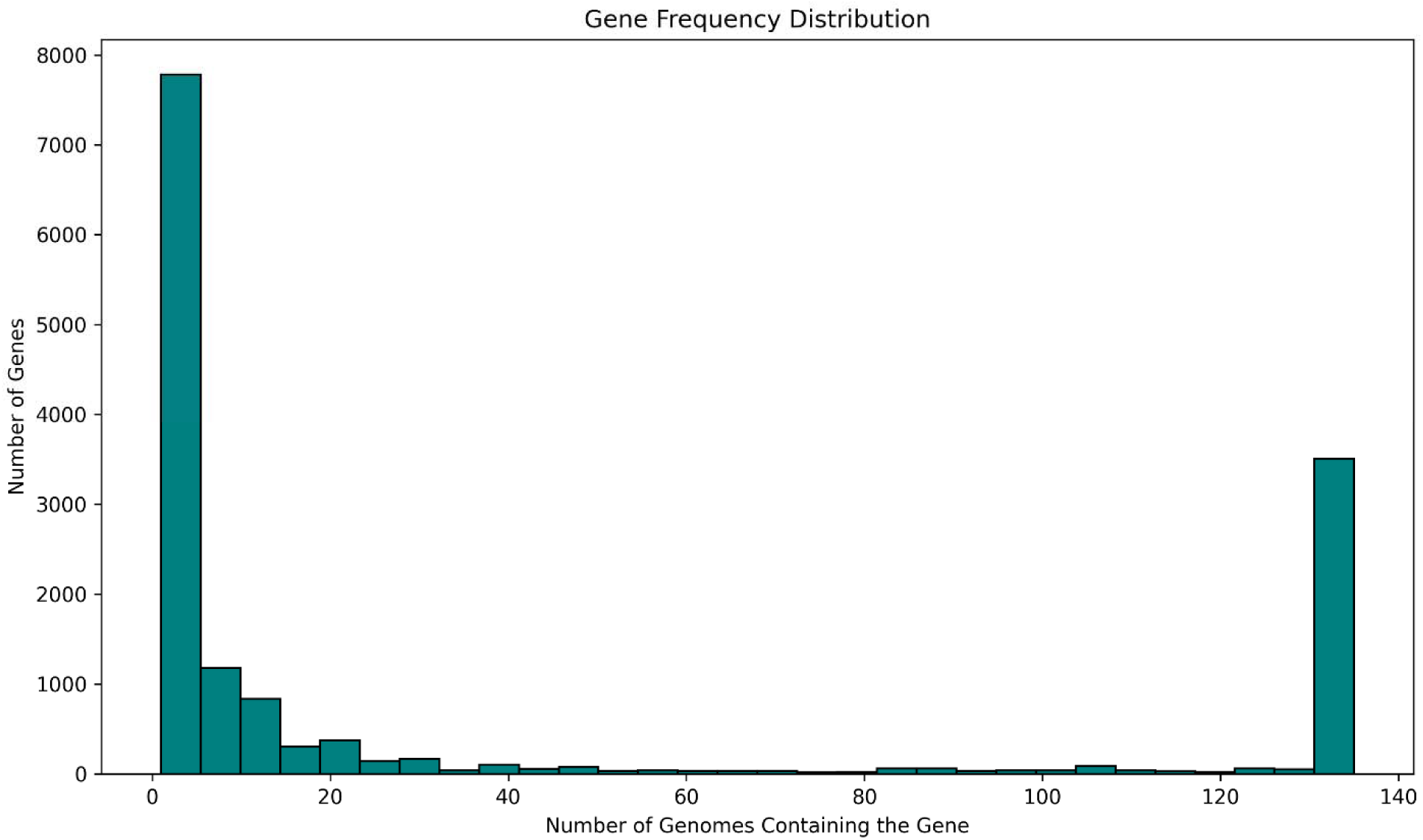
Gene frequency distribution among 135 *Salmonella enterica* genomes. The plot illustrates the distribution of gene presence across the analyzed genomes, with the x-axis representing the number of genomes in which a given gene is present and the y-axis representing the number of such genes. A bimodal distribution is observed, with peaks at the extreme left (representing rare and strain-specific genes) and right (ubiquitous, core genes). This points to the open nature of the *Salmonella* pan-genome and the coexistence of highly conserved and highly variable gene subsets.

### 3.3 AMR Resistance Genes

Genome mining of virulence genes revealed that *Salmonella* isolates carry a wide array of antimicrobial resistance genes across almost all major drug classes. β-lactamase genes appeared frequently, ranging from narrow-spectrum TEM variants to broader-spectrum enzymes such as *blaCMY*-2 and extended-spectrum β-lactamases including *blaCTX*-M-2, *blaCTX*-M-9, and *blaCTX*-M-55. The detection of *blaOXA*-232 is concerning because it can confer resistance to carbapenems, which are considered a last-resort treatment. Several isolates also carried plasmid-mediated quinolone resistance genes, including *qnrB19* and *qnrS1*, such that the administration of fluoroquinolones against the isolates would be ineffective. Genes conferring resistance to trimethoprim-sulfamethoxazole, such as multiple *sul* and *dfrA* variants were widespread. The prevalence of trimethoprim-sulfamethoxazole resistance genes is worrying because these drugs are often preferred for first-line therapy against common infections (Sivanandy et al., 2025). We also noted that aminoglycoside-modifying enzymes, tetracycline efflux pumps (tet[A–C, G]), macrolide inactivation (*ereA*), and chloramphenicol/florfenicol resistance genes (*catA1, floR*) appeared in many strains. The presence of macrolide inactivation genes (*ereA*) can render *Salmonella* resistant to azithromycin, which is a critical treatment for infections in medicine and agriculture (Zieliński et al., 2021). This would mean the use of last-resort antibiotics and a growing cycle of multidrug resistance. Of particular concern, some genomes carried *mcr*-*1.1* and *mcr-9*, which provide transferable colistin resistance, and *fosA7*, which inactivates fosfomycin. The presence of these genes highlights the potential for multidrug resistance to spread between humans, animals, and the environment. Clinically, these findings are worrisome as they threaten the effectiveness of key drugs used in salmonellosis treatment, especially the third-generation cephalosporins, fluoroquinolones, and carbapenems, which are used in rare but severe cases. Figure 6 shows the frequencies of the top 20 AMR genes across various genomes. Among the 135 AMR-positive *Salmonella* genomes analyzed, the most common genes were *aac(6’)-laa_1* (n=134), *tet(A)_6* (n=59), *fosA71* (n=34), *sul2_2* (n=32), *alph(6)-ld_1* (n=30), *sul_5 (n=29)*, *aaph(3’)-la_1* (n=18), *qnrB19_1* (n=18), *ant(3")-la_1* (n=18), *blaTEM-1B_1* (n= 17). The most prevalent antimicrobial resistance gene was *aac(6’)-laa_1*, detected in nearly all isolates (99%), suggesting its potential role as a conserved or chromosomally encoded resistance determinant. The AMR gene profile reveals a predominance of aminoglycoside resistance genes and widespread baseline resistance to this antibiotic class. The tetracycline resistance gene *tet(A)_6* was the second most frequent, found in 59 genomes (43.3%), followed by *fosA7_1* and *aph(6)-Id_1*, each present in at least 30 genomes. Sulfonamide resistance genes (*sul1_5* and *sul2_2*) were also commonly encountered, highlighting widespread resistance to sulfonamides. Notably, *blaTEM-1B_1*, a β-lactamase gene associated with penicillin resistance, was detected in 17 isolates, and *qnrB19*_*1*, conferring quinolone resistance, was present in 18 genomes. The presence of multiple aminoglycoside-modifying enzymes, including *ant(3’’)-la_1* and *aph(3’)-la_1*, further illustrates the diverse resistance gene landscape in *Salmonella* isolates circulating in the African food chain. Overall, our findings suggest that the co-occurrence of resistance to tetracyclines, fosfomycin, sulfonamides, and beta-lactams, demonstrating the multidrug-resistant nature of the *Salmonella* population under study.

**Figure 6:**
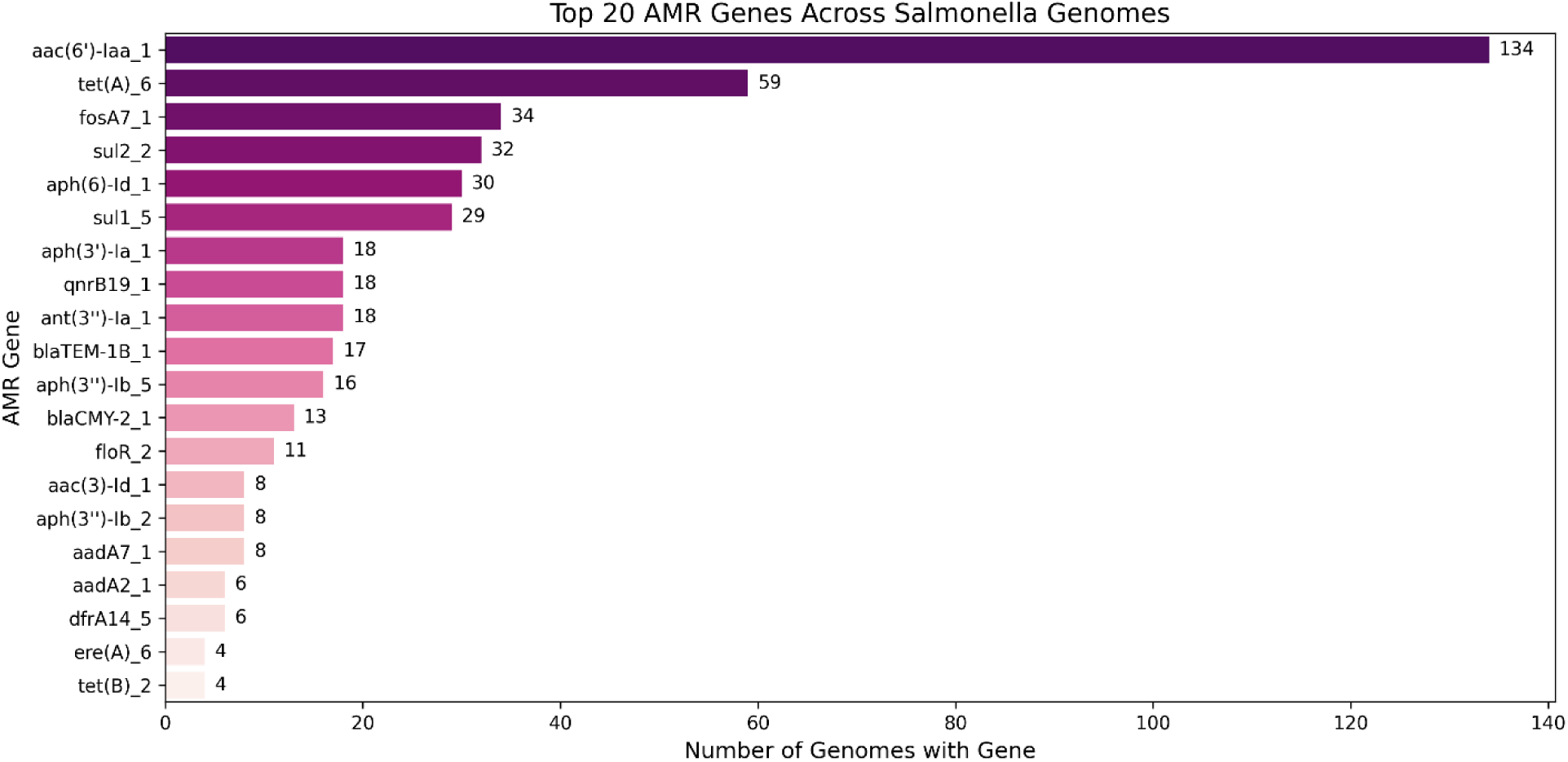
Distribution of the 20 Most Prevalent AMR Genes Among 135 AMR-Positive *Salmonella enterica* Genomes

Figure 7 below shows a binary heatmap depicting the presence and absence of AMR genes across *Salmonella* genomes, revealing distinct patterns of resistance distribution. The aminoglycoside resistance gene *aac(6’)-laa_1* was universally present, appearing across all genomes. This is likely an indication of the potential chromosomal origin or lineage fixation of this gene. In contrast, other genes such as *tet(A)_6*, *fosA7_1*, *aph(6)-ld_1*, and *sul1_5* exhibited heterogeneous distribution, with clusters of genomes harboring multiple resistance determinants. The heatmap also highlighted varying degrees of AMR gene co-occurrence, particularly among tetracycline, sulfonamide, and aminoglycoside genes. This genomic diversity in resistance profiles is indicative of potential the role of horizontal gene transfer and niche adaptation in shaping the AMR landscape within foodborne *Salmonella* strains in Africa.

**Figure 7:**
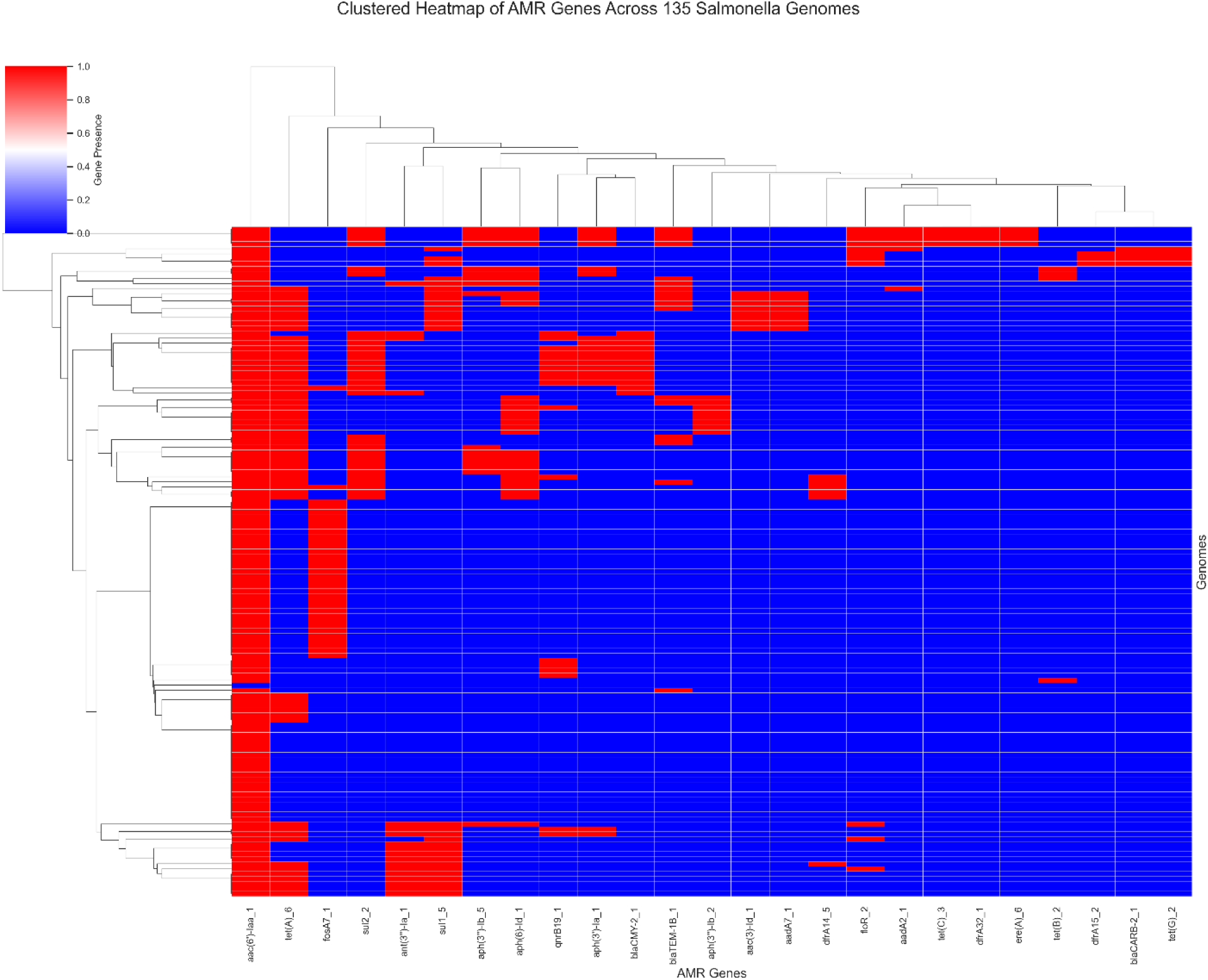
Presence–Absence Matrix of Antimicrobial Resistance Genes in AMR-Positive *Salmonella enterica* Genomes from African food chains

### 3.4 Structure and Distribution of AMR Genes

We used the Shannon diversity index to quantify AMR gene richness and evenness within individual genomes and across ecological and taxonomic strata. Fig. 8 below shows the distribution of these genes across various variables. It is evident that considerable inter-isolate variability exists whereby some genomes harbor highly diverse resistance repertoires. When categorized by source type (Fig. 8B), isolates from avian, poultry, and unspecified mean origins exhibited the highest AMR diversity, consistent with known antimicrobial practices in livestock systems. Serovar-based analysis (Fig. 8C) showed that Heidelberg and Enteritidis carried more complex resistance profiles, which is likely due to intrinsic lineage traits or extrinsic ecological exposure. Regional variation (Fig. 8D) revealed that isolates from Burkina Faso, South Africa, Tanzania, and Tunisia had greater diversity and variation in AMR gene content. This could be an indication of a heterogeneous AMR gene pool, likely due to different sources of contamination, selective pressures, or diverse clonal lineages in these countries. These countries are potential regional selection hotspots. These findings highlight the utility of integrating ecological metadata with genomic surveillance to identify high-risk reservoir and inform AMR mitigation strategies.

**Figure 8.**
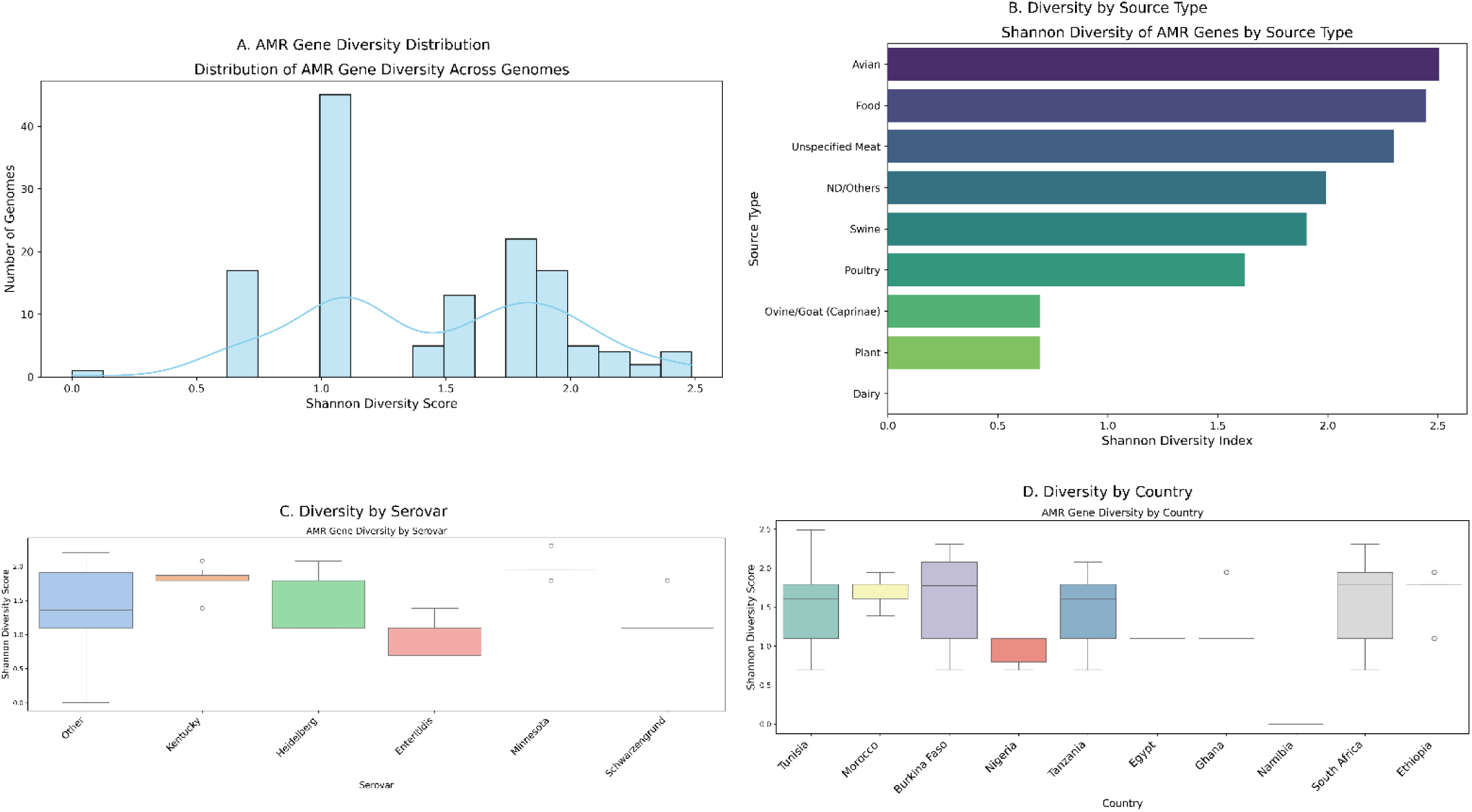
Shannon diversity of AMR genes across the variables (A) AMR gene diversity per isolate. (B) Distribution by source type. (C) Stratification by serovar, and (D) Regional variation by country of isolation.

We used NetworkX (https://github.com/networkx/networkx) for analysis of co-occurrence of antimicrobial resistance (AMR) genes whereby our results revealed structured and non-random patterns of association. These patterns could be suggestive of mobile genetic elements and horizontal gene transfer mechanisms. Our findings point to the co-occurrence of aminoglycoside, beta-lactam, tetracycline, and quinolone resistance genes in *Salmonella* isolates. In the network view (Fig. 9-A), highly connected hubs such as *aac(6’)-laa_1*, *tet(A)_6*, and *sul2_2* emerged as central nodes, often co-localizing within the same genome. These genes were linked to other major nodes include *ant(3”)-la_1, qnrB19_1, blaCMY-2_1, aadA*, and *aph(3’)-la_1, aph(3’’)-lb_5,* and *aph(6)-ld_1.* These associations suggest the presence of multidrug resistance cassettes and/or integrons. The corresponding matrix (Fig. 9-B) illustrates that several gene pairs frequently co-occurred across multiple isolates, reinforcing the hypothesis of coordinated dissemination. The dense connectivity among these genes depicts the modular nature of AMR gene acquisition and the potential role of mobile genetic elements, such as plasmids and transposons in facilitating their spread. The conserved nature of these resistance modules across diverse serovars and sources points to common selective pressures within African food production environments. This understanding may be important in identifying priority resistance loci for molecular surveillance and targeting potential vectors of horizontal gene transfer.

**Figure 9.**
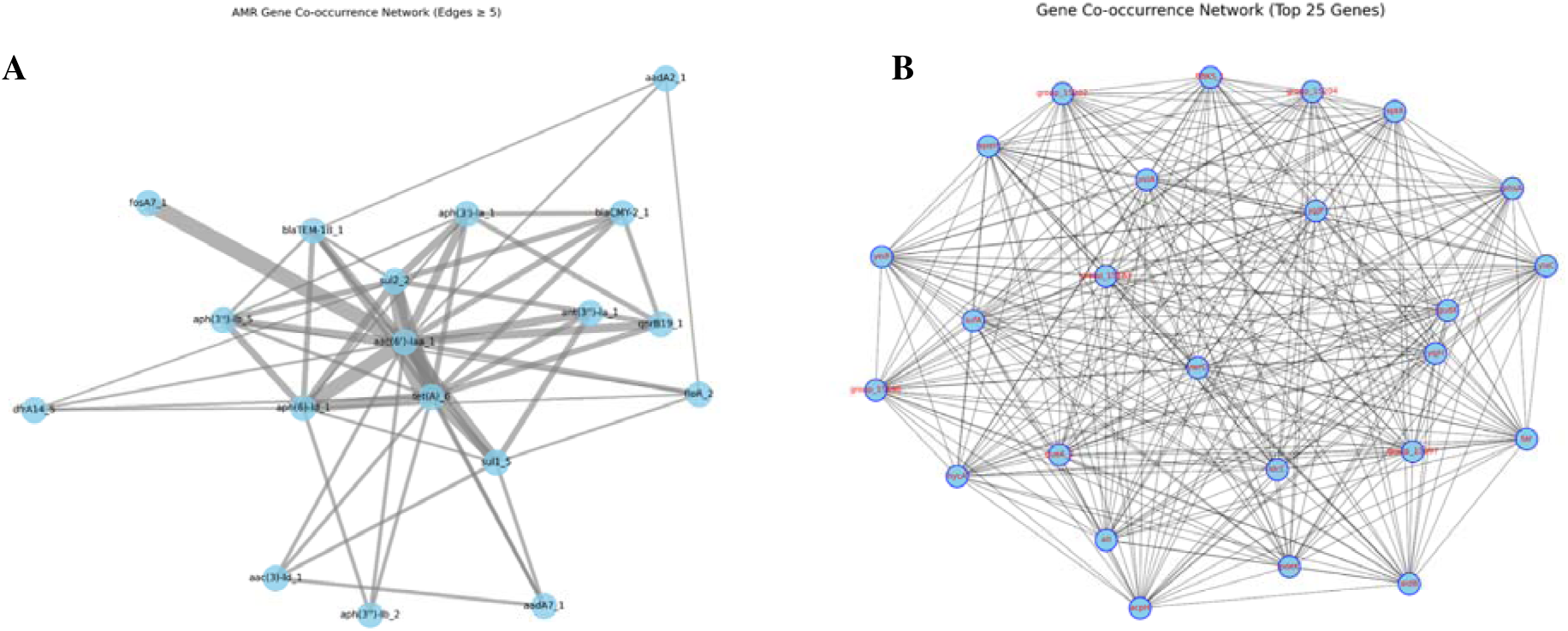
Co-occurrence patterns of AMR genes in Salmonella enterica. Fig.9(A) shows AMR gene co-occurrence *network* of interactions among frequently detected resistance genes. Fig. 9(B) is the Matrix view of the top 25 most frequent AMR gene co-occurrence patterns across isolates.

### 3.5 Plasmid burden and virulence gene presence

We analyzed plasmid burden and virulence gene content as components of the accessory genome with the results showing significant heterogeneity across serovars. Fig. 10 shows that isolates belonging to Muenchen, Heidelberg, Infantis, and *Typhimurium* serovars displayed higher plasmid counts than the rest. This could be indicative of enhanced genomic plasticity and a greater propensity for acquiring resistance elements. These serovars are often associated with broad host ranges and zoonotic transmission. As a result, plasmid-mediated traits likely facilitate adaptation to diverse ecological niches. Fig. 11 presents a heatmap of key virulence genes across various assemblies. Genes like *csgB*, *csgE*, *fimD*, *invA*, *invE*, *invH*, *mig-14*, *orgA*, *pipB2*, *sinP*, *sinH*, *sopD*, *spaO*, *spaR*, *SSaE*, *ssaI*, *ssaL*, *ssaR*, *ssaU*, *sscB*, *sseC*, and *steA* are nearly universally present in the *Salmonella* samples. It is likely that genes are part of the core virulence genome of *Salmonella enterica* among the foodborne serovars. Their universal presence suggests that the pathogenic potential is conserved even across genetically diverse lineages. These genes are likely components of the Type III Secretion Systems (T3SS) located on *Salmonella* Pathogenicity Islands (SPIs) such as SPI-1 and SPI-2, which are essential systems of host cell invasion, intracellular survival, and systemic spread as previously reported (Hansen-Wester and Hensel, 2001; Miki et al., 2009; Yu et al., 2010). Again *csg* and *film* cluster genes like *fimD* are well known mediators of biofilm production and related virulence in *Salmonella* (Musa et al., 2024). On the other hand, genes such as *papB*, *papE*, *papH*, *papK*, *pefC*, *rcK*, *tcpC*, *ybtP*, and *ybtT* were rare across samples and serovars. The rarity of these genes may point to specificity of ecological niches rather than accessory virulence. Nonetheless, the diversity in virulence profiles points to lineage-specific pathogenic mechanisms. Our results are suggestive of accessory genome’s role in shaping virulence in *Salmonella*.

**Figure 10.**
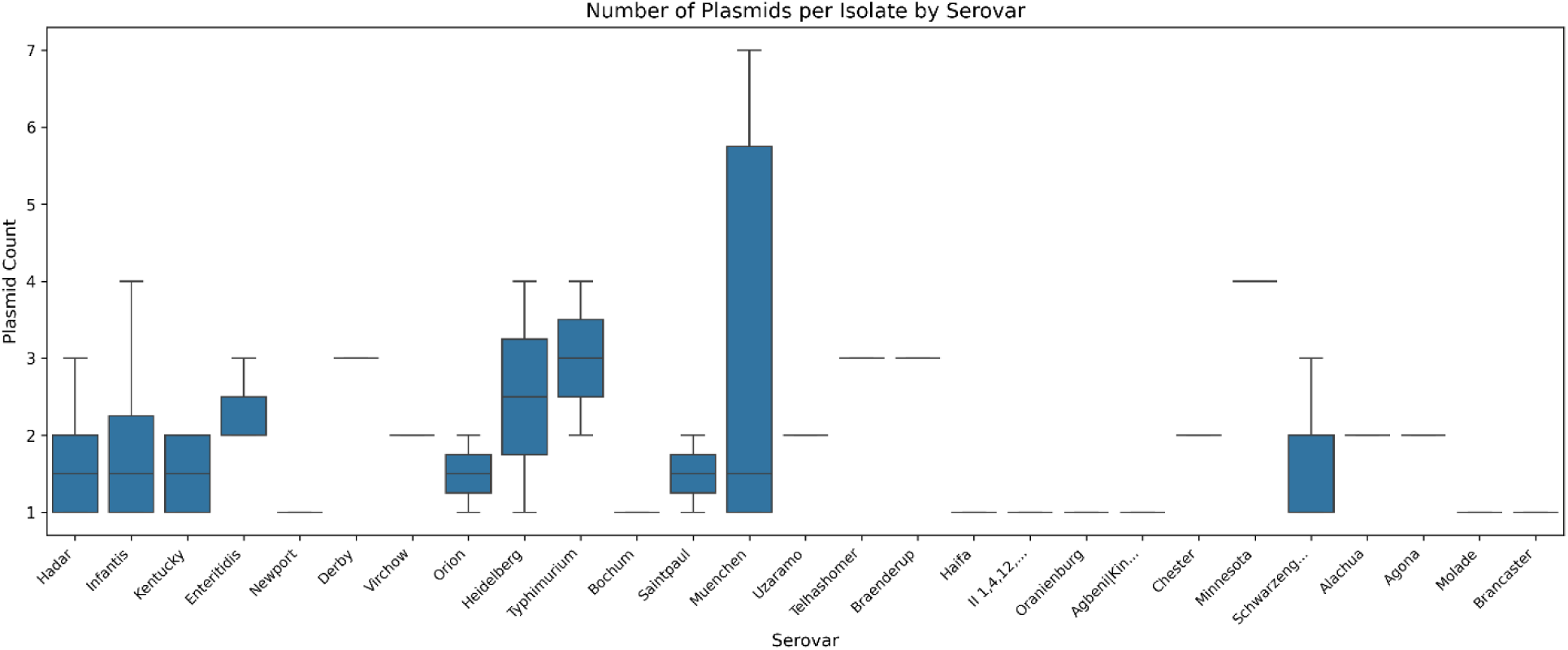
: A boxplot showing the number of plasmids per isolate by serovar

**Figure 11:**
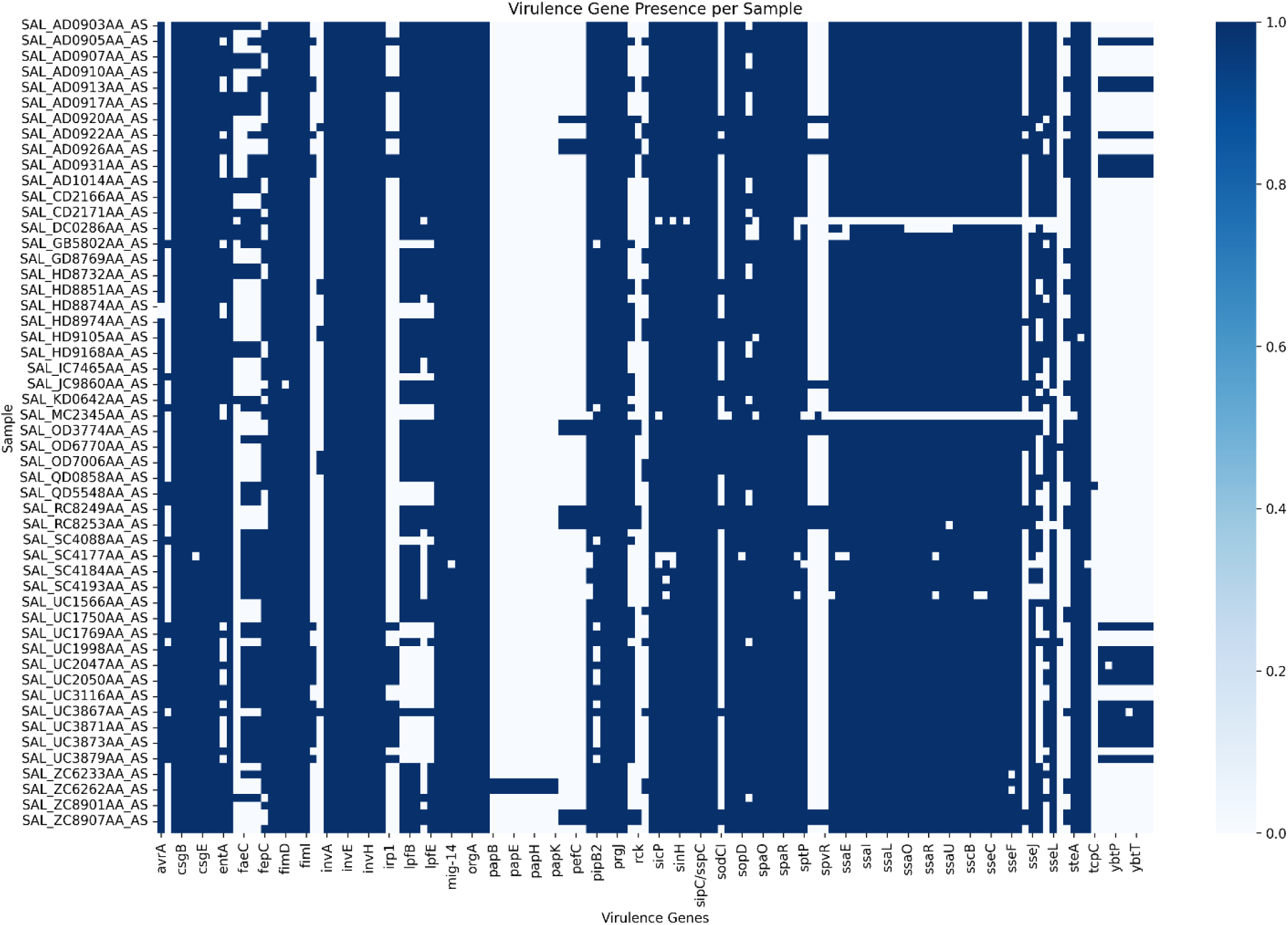
A heatmap depicting the distribution of virulence genes in AMR positive *Salmonella genomes.* The binary heatmap illustrates the presence (dark blue) and absence (light blue) of key virulence genes across 135 *Salmonella enterica* genomes carrying antimicrobial resistance (AMR) determinants. Each row represents an individual genome while each column corresponds to a virulence gene.

### 3.6. Quorum sensing and biofilm genes

We then conducted a manual curation of quorum sensing and biofilm-associated genes. For this, we searched the gene presence and absence data for presence of common Salmonella quorum sensing genes as identified in literature. The distribution of biofilm and quorum-sensing genes across the Salmonella isolates are shown in Figure 12. The figure shows a conserved set of regulatory and structural components that support biofilm formation, cell-to-cell communication, and adaptation to stress. Classical biofilm genes, including *csgA*, *csgB*, *csgC*, *csgD*, and *csgG*, which are involved in curli assembly and regulation, appeared in nearly all 135 isolates. Genes for cellulose synthesis, such as *bcsA*, *bcsB*, and *bcsC*, were also widely present. Together, curli and cellulose form the main components of the Salmonella extracellular matrix. The isolates also carried *bdcA*, a protein that binds cyclic-di-GMP and controls biofilm dispersal, allowing the bacteria to switch between free-living and surface-attached states depending on environmental conditions. Quorum-sensing genes were common across the isolates. AI-2 signaling system genes, including *luxS*, *lsrA*, *lsrB*, *lsrC*, *lsrD*, and *lsrK*, were found in more than 90 percent of genomes, which is a pointer to a conserved role for interspecies communication. A single quorum-quenching gene, *aidA*, appeared in 22 isolates. Its presence may allow these strains to interfere with the quorum-sensing behaviors of other bacteria, reducing biofilm formation or affecting virulence in mixed communities. This could give Salmonella an edge in environments such as the gut or on food-processing surfaces. Motility and flagellar regulatory genes, including *flhC*, *flhD*, *ydiV_1*, and *ydiV_2*, were also widespread. The fact that most isolates carried these genes, show that motility and flagellar regulation are important for initial biofilm formation. The frequent co-occurrence of motility and biofilm genes suggests that *Salmonella* coordinates surface attachment, biofilm development, and dispersal to adapt to different environments.

**Figure 12.**
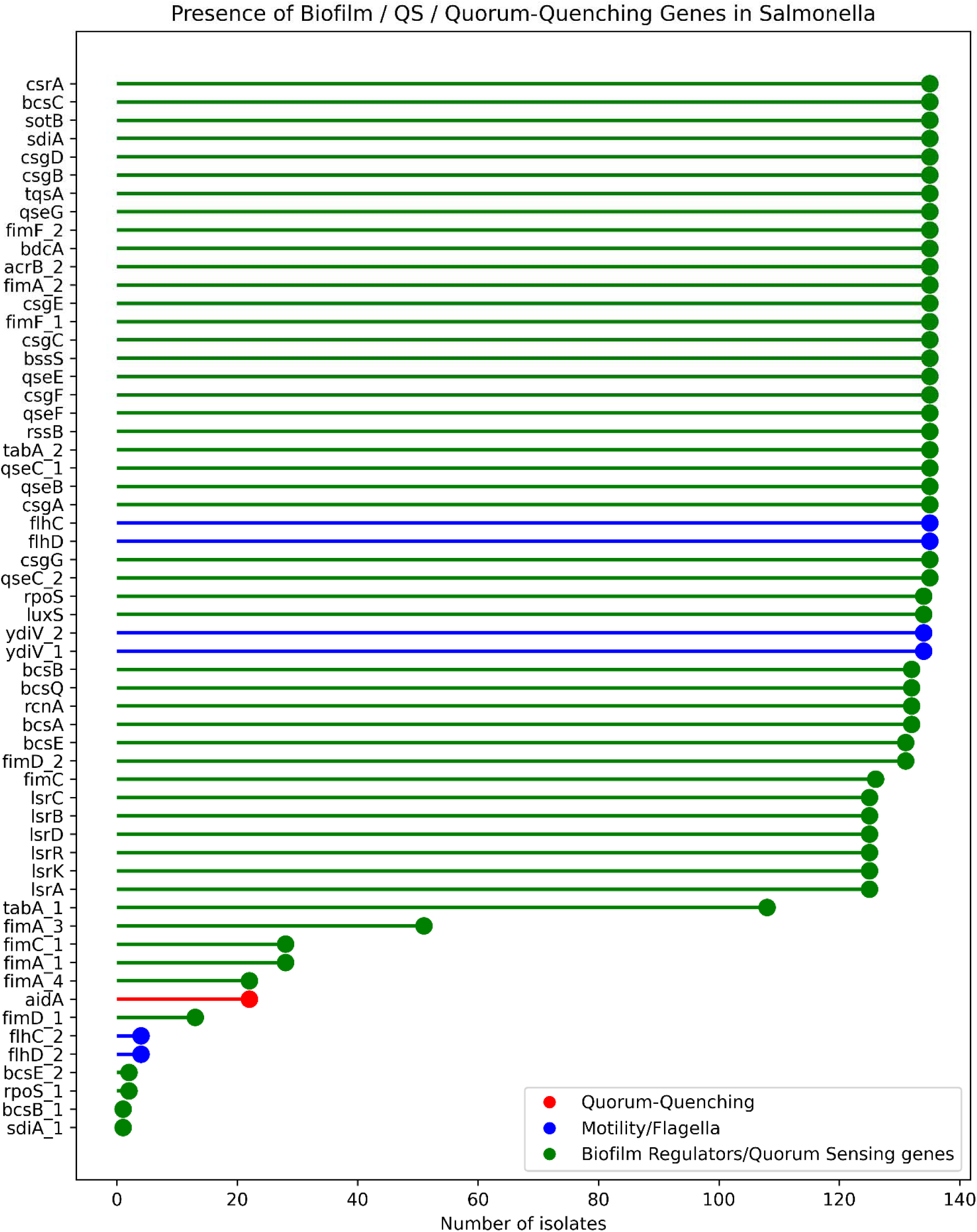
A plot showing the presence of biofilm- and quorum-sensing–related genes across Salmonella isolates. Each dot represents the number of isolates in which a gene was detected. Lines and markers are color-coded by functional category: quorum-quenching gene (red), motility/flagellar regulators (blue), and other biofilm or quorum-sensing genes (green).

## 4. Discussion

This study involved the analysis of genomic data of *Salmonella* isolates from food samples in Africa. Phylogenetic analysis using cgMLST-based HierCC clustering revealed tight, monophyletic clades at HC100, reflecting high genetic relatedness. Tight clustering at HC100 could be indicative of the presence of long-term endemic persistence (Zhou et al., 2020). The clustering also points to the possibility of the isolates sharing a high number of core genomes MLST alleles. This hypothesis is supported by the presence of a large number of core genome of 3,350 genomes present across the isolates. Broader clustering at HC2000 uncovered super-lineages that grouped multiple serovars, supporting the notion of shared ancestral backgrounds and highlighting the limitations of serotyping in capturing true evolutionary relationships. Lineage-specific gene association testing further identified genetic markers exclusive to certain serovars, opening avenues for targeted diagnostics and surveillance.

We also conducted a comprehensive pan-genomic and AMR landscape analysis of *Salmonella enterica* strains from African food chains. Our analysis focused on the distribution of resistance genes and their relationship to genetic lineage and phylogenetic structure. After analyzing 135 AMR-positive *Salmonella* genomes from cross the continent, we identified a total of 15,269 genes constituting the pangenome. Of these, 3,350 were core genes shared across nearly all isolates (99–100%), while the cloud genome representing rare or unique genes accounted for the largest proportion with 10,295 genes. These figures reflect a moderately closed pangenome structure with but with considerable genomic fluidity within the population, in line with earlier reports (Chand et al., 2020). The γ parameter of 0.2844 further supports the closed genome nature of the *Salmonella* isolates from African food chains. Our findings are consistent with earlier reports of a closed genome (γ = 0.235909) in *Salmonella enterica* serovar (Coluzzi et al., 2025). Thus, the identification of new genomic regions in future will decline for each new genome (Laing et al., 2017; Tettelin et al., 2005).

AMR profiling revealed that the aminoglycoside resistance gene *aac(6’)-laa_1* was nearly universal, occurring in 99% of isolates. This is consistent with an earlier study that reported the presence of aminoglycoside resistance genes in nearly all food isolates (Pettengill et al., 2020). Resistance to aminoglycosides occurs due to mutations that result in formation of aminoglycoside-modifying enzymes (AMEs) whose primary mechanism is the prevention of the antibiotic from binding the bacterial target (Kumar et al., 2025). Tetracycline (tetA), sulfonamide (sul2), and fosfomycin (fosA71) resistance genes were also widely distributed, indicating a multidrug-resistant (MDR) population structure. Previous studies in literature have shown the predominance of these genes in *Salmonella* isolates (Pavelquesi et al., 2021). Genes such as *qnrB19_1*, *ant(3”)-la_1*, and *blaTEM-1B_1* were frequently co-localized in isolates, suggesting the presence of mobile genetic elements like plasmids or genomic islands that serve as vectors for AMR gene dissemination. This is further supported by the prominence of co-occurrence networks where hub genes such as *aac(6’)-laa_1*, *tet(A)_6*, and *sul2_2* formed central nodes with multiple connections, indicating a potential for coordinated gene mobility and persistence.

We further established that core virulence determinants such as *invA*, *ssaL*, *ssaU*, *spaO*, and *steA* were consistently present across isolates and serovars. This could mean the presence of a conserved pathogenic backbone that is essential for *Salmonella* infection and intracellular survival. The near-universal presence of these genes highlights their evolutionary conservation and indispensability for host invasion, aligning with previous findings that place the Type III secretion system (T3SS) and associated SPI-1/SPI-2 effectors at the core of *Salmonella* pathogenicity (Jennings et al., 2017; Miki et al., 2009). *Salmonella* pathogenicity island-1 (SPI-1) are present in *Salmonella* lineages and have the role of encoding a type III secretion system (TTSS-1) that is considered essential for invasion of intestinal epithelial cells, host adaptation and survival in the gastrointestinal tract (Lou et al., 2019; Phoebe Lostroh and Lee, 2001). Conversely, rare virulence genes such as papE, rcK, and tcpC were infrequently detected, reflecting their lineage-specific acquisition or possible roles in niche adaptation.

Diversity analyses using Shannon indices indicated significant variation in AMR gene richness across geographical locations. For example, the isolates from Tunisia showed high interquartile variability and extended error bars, which may be an indication of heterogeneous AMR acquisition. Previous studies from Tunisia have shown the presence of multi-drug resistance *Salmonella* (Oueslati et al., 2023). Similarly, higher diversity indices were noted for Burkina Faso, South Africa, Tanzania. This aligns with previous studies showing diverse *Salmonella* serotypes and antimicrobial resistance in sub-Saharan countries (Kilongosi Webale, 2024; Mwambene et al., 2025; Ramatla et al., 2021; Siourimè et al., 2017). In contrast, countries like Morocco displayed more stable profiles, indicating a potentially localized resistance gene pool. Such diversity underlines the importance of regional surveillance and suggests that resistance dynamics may not be uniformly distributed across Africa. Overall, genome analysis points to genomic complexity and a high AMR burden of *Salmonella* in African food chains. The detection of lineage-specific accessory genes, core virulence traits, and diverse AMR determinants supports the need for integrated genomic surveillance to guide antimicrobial stewardship and public health interventions.

## 5. Conclusion

This study provides a comprehensive analysis of AMR-positive *Salmonella enterica* genomes from African food chains. The findings show important trends in antimicrobial resistance, virulence, and lineage-specific gene content. While having a moderately closed pangenome, *Salmonella* isolates from African food chains show possibility of flexible gene acquisition. Hierarchical clustering confirmed the presence of serovar- and lineage-associated genes, suggesting potential targets for molecular surveillance and a conservation of core virulence determinants. Furthermore, country-level differences in AMR gene diversity highlight the impact of local selection pressures and the need for region-specific mitigation strategies. These findings show the value of genome-wide approaches in advancing our understanding of *Salmonella* epidemiology. The findings also attest to the need for the integration of genomic data into public health and food safety monitoring across Africa.

